# Increased *in vivo* transduction of AAV-9 cargo in Alport podocytes

**DOI:** 10.1101/2025.04.28.650965

**Authors:** Emily Williams, Maryline Fresquet, Gema Bolas, Shota Kaseda, Kevin A. Goncalves, Andrew Steinsapir, Antony Adamson, David R Sherwood, Rachel Lennon

**Affiliations:** Manchester Cell-Matrix Centre, Division of Cell-Matrix Biology and Regenerative Medicine, School of Biological Sciences, Faculty of Biology Medicine and Health, The University of Manchester, Manchester Academic Health Science Centre, Manchester, M13 9PT, UK; Deerfield Discovery and Development, Deerfield Management Company, New York, NY, USA; Genome Editing Unit Core Facility, Faculty of Biology, Medicine and Health, University of Manchester, Manchester M13 9PT, UK; Department of Biology, Duke University, Box 90338, Durham, NC 27708, USA; Department of Pediatric Nephrology, Royal Manchester Children’s Hospital, Manchester University Hospitals NHS Foundation Trust, Manchester Academic Health Science Centre, Manchester, M13 9WL, UK

**Keywords:** Alport syndrome, gene therapy, AAV, glomerulus, podocyte, collagen IV

## Abstract

Alport syndrome is a rare genetic disorder characterized by progressive kidney disease, hearing loss, and eye abnormalities. Gene therapy for Alport syndrome has not yet been realized due to technical challenges, including effective transduction of target cells. In this study, we established a quantitative testing platform using podocytes in culture, *ex vivo* glomeruli, and a mouse model of Alport syndrome to evaluate the transduction efficacy of adeno-associated Virus-9 (AAV9) as a gene delivery vehicle. We compared transduction levels of AAV9-GFP vectors between healthy and Alport podocytes in culture, revealing that both cell types exhibited similar transduction rates. We then incubated *ex vivo* glomeruli with AAV9-GFP and found enhanced transduction in Alport compared to wild type podocytes. Finally in mice following a peripheral intravenous injection of AAV9-GFP we found a striking increase in transduction in Alport podocytes suggesting that the pathological environment may facilitate higher penetration of the vector. These findings underscore the potential of AAV9 for effective gene delivery in the context of Alport syndrome, providing a foundation for future therapeutic strategies aimed at correcting the underlying genetic defects.

**Plain Language Summary:** This study highlights the clinical significance of AAV9 as a potential gene delivery vehicle for Alport syndrome treatment. Our findings demonstrate that Alport podocytes exhibit enhanced transduction levels in *ex vivo* and *in vivo* environments compared to healthy podocytes. This suggests that AAV9 could effectively target podocytes in Alport syndrome, paving the way for innovative gene therapy strategies that may improve patient outcomes and quality of life.

## Introduction

Alport syndrome is a genetic disorder characterized by progressive kidney disease, hearing loss, and eye abnormalities. It is caused by variants in the genes *COL4A3, COL4A4*, or *COL4A5*, which play a crucial role in the assembly of type IV collagen, a vital component of basement membranes (1). Defects in collagen IV assembly, particularly the α345(IV) scaffold, can disrupt the glomerular basement membrane (GBM) and impair its filtration function (2-4). Despite the identification of its genetic underpinnings over 30 years ago, advances in gene therapy have been slow due to challenges in delivering genes specifically to the kidneys and the large size of the collagen genes, which exceed the cargo capacity of many commonly used vectors (5). Since podocytes are the cellular source of the collagen α345(IV) network affected in Alport syndrome (6), this condition is regarded as a podocytopathy. Consequently, the podocyte is the main target cell for genetic therapies in Alport syndrome (7).

Recent developments in genetic engineering have introduced a diverse range of vectors designed to deliver genetic cargo to cells for therapeutic purposes, including viral and non-viral options such as plasmids, liposomes, and nanoparticles (8, 9). One promising vehicle is adeno-associated virus (AAV), a non-pathogenic viral vector frequently used in gene therapy that exhibits a low risk of adverse effects in humans (10). AAV can enter cells through receptor-mediated endocytosis. Following endosomal escape, the vector is transported to the nucleus, where it releases its genetic material. Host cell machinery is then used for gene expression (11). This property allows AAV vectors to induce stable, long-term gene expression, offering potential benefits for chronic kidney diseases like Alport syndrome (12) and for transduction of terminally differentiated cells such as podocytes.

The selection of an AAV capsid and the delivery method have been shown to influence the kidney cell types that can be targeted for gene therapy (5). Naturally occurring AAV serotypes have varying levels of tropism for kidney tissue with AAV2, AAV4, AAV8, AAV9, AAV10, and AAV11 all displaying affinity for the kidney (13). However, previous studies found AAV9 to be the most effective capsid for kidney cell transduction across various administration routes. Intravenous administration of AAV9 primarily results in the transduction of mesangial and interstitial cells in healthy kidneys (14, 15). While in a chronic kidney disease mouse model, intravenous injection of AAV9 showed effective transduction of podocytes and proximal tubules (16).

This study aimed to evaluate the transduction efficacy of AAV9 with a green fluorescent protein cargo (ssAAV9-GFP) in healthy and Alport podocytes via intravenous injection. We tested increasing doses of ssAAV9-GFP in 8-week-old mice and its effect during disease progression in 8-, 10- and 12-week-old mice. To facilitate this assessment, we established a testing platform comprising mouse and human podocytes, *ex vivo* glomeruli, and a murine model of Alport syndrome. This platform enables quantitative evaluation of podocyte transduction and supports the advancement of gene therapies for glomerular disorders, including Alport syndrome.

## Methods

### AAV9-GFP production

The AAV9-GFP virions were produced at Viralgen (Spain) in Pro10™ cells by triple transfection with the GFP (green fluorescent protein) transgene under the CMV promoter (pALD-ITR-WPRE-GFP, Aldevron, 5069-10, lot 132222, **S. Figure 1**), a plasmid carrying pAAVrep2capX and a helper plasmid carrying the VA-RNA, E2A and E4 helper genes of the Adenovirus serotype 9. Material from the bioreactor was clarified by depth filtration prior to the first purification step by immunoaffinity chromatography. The pre-purified product was then subjected to a DNA-containing AAV “full” particles enrichment step using iodixanol gradient ultracentrifugation prior to a second chromatography step using an ion exchange matrix. The vector was produced using preclinical research standards that met criteria for in vivo research use in terms of percentage of empty capsids, total purity and endotoxin levels. The vector genome titer (3.3×10^14^ vg/mL) and purity of the resulting single-stranded adeno-associated virus ssAAV9-GFP were confirmed by ITR-ddPCR and SDS-PAGE protein gel analysis.

### Animal model

*Col4a5* knockout (KO) mice, an X-linked Alport model, were obtained from the International Mouse Phenotyping Consortium (17). The line was generated by deletion of the critical exon 36, resulting in the absence of the collagen-α5(IV) protein, using the published allele map (18). These mice were bred on a C57Bl/6 background and were previously characterized (19). Hemizygous male mice were selected for phenotypic consistency and wild type (WT) littermates were used as controls. 3-4 mice per group/condition were injected with AAV9 for transduction efficiency analysis. Further details on the groups, age of the mice, dose injected can be found in the “AAV9 transduction” section. Mice were housed at The University of Manchester Animal Unit under Specific Pathogen Free (SPF) conditions. All experiments were approved by The University of Manchester Local Review Committee and conducted in compliance with UK Home Office Animals (Scientific Procedures) Act 1986 regulations and in accordance with the ARRIVE guidelines.

### Glomerular isolation and cell culture

Cortical tissue was dissected from fresh mouse kidney and cut into small pieces (≤1 mm^3^). Tissue pieces were pressed through a 100µm cell strainer (Appleton Scientific, Cat #ACF963) and washed with sterile PBS. The crude isolate was then passed through a 70μm cell strainer (Appleton Scientific, ACF963), where glomeruli were captured. Glomeruli were washed with PBS and seeded onto a 100 mm cell culture dish. Cells were cultured in RPMI 1640 medium with 10% FBS (Gibco, Cat# A-5670801), 1% insulin-transferrin-selenite (ThermoFisher, Cat# 1400045) and 1% Penicillin-Streptomycin (Sigma-Aldrich, Cat# 15070063). After 14 days of culturing, primary mouse podocytes outgrowths were detected and used for the multiplicity of infection (MOI) assay.

Human kidney-derived podocytes cell line PODO/TERT256 (Evercyte, CHT-033-0256; RRID:CVCL_JL76) were cultured in PodoUp3 medium (Evercyte, MHT-033-3) and used for the Multiplicity of Infection (MOI) assay.

### AAV9 transduction

A single 100μL aliquot of AAV9-GFP was thawed from −80°C and stored at 4°C. AAV9-GFP was diluted in cell culture media to the desired concentration for *in vitro* assays or in sterile diluent for *in vivo* injections.

#### In cultured podocytes

For cell culture experiments, 1.6×10^5^ PODO/TERT256 (RRID:CVCL_JL76) and isolated primary podocytes from wild type and Alport mice were plated on laminin-coated cover slips in a 24 well plate. Serial dilutions of AAV9-GFP were performed to achieve an MOI of 5×10^4^ to 3.125×10^7^. Cells were incubated with AAV9-GFP for 24 hours. Cells were fixed with 4% paraformaldehyde for 15 minutes, washed with PBS and blocked with 5% BSA, 0.3% Triton-X in PBS for 1 hour at room temperature. After blocking, cells were washed with PBS and incubated with anti-GFP (1:500, Abcam Cat# ab6673; RRID:AB_305643) and primary podocytes with anti-WT1 antibody (1:300, Abcam Cat# ab89901; RRID:AB_2043201) in PBS 3% BSA, 0.1% Tween overnight at 4°C. The next day cells were incubated with donkey anti-rabbit Alexa Fluor 647 (1:1000, Thermo Fisher Scientific Cat# A-31573; RRID:AB_2536183) and anti-goat Alexa Fluor 555 (1:1000, Abcam Cat# ab150130; RRID:AB_2927775) for 1 hour at room temperature. DAPI (1:2000, Cell Signaling 4083) was used for nuclear staining. Images were collected on a Zeiss Axioimager.D2 upright microscope using a 20x / 0.5 EC Plan-neofluar objective and captured using a Coolsnap HQ2 camera (Photometrics) through Micromanager software v1.4.23. Images were then processed and analyzed using Fiji ImageJ.

#### In isolated mouse glomeruli

For *ex vivo* experiments, isolation of glomeruli from 8-week-old WT and Col4a5 KO Alport mice was performed as previously described. Briefly, glomeruli captured in the 70μm cell strainer were seeded onto a 24 well dish with 8.8×10^11^ vg/mL of AAV9-GFP and incubated for 48 hours. After viral transduction, glomeruli were centrifuged at 400xg for 5 minutes, washed in PBS and fixed with 2% paraformaldehyde for 15 minutes. Glomeruli were permeabilized and blocked in 3% BSA, 0.1% Tween, 0.2% Triton X-100 for 1 hour at room temperature. Glomeruli were then incubated with anti-GFP and anti-WT1 antibodies (1:250, Abcam Cat# ab6673; RRID:AB_305643, Abcam Cat# ab89901; RRID:AB_2043) in blocking buffer overnight at 4°C. The next day glomeruli were incubated with donkey anti-goat Alexa Fluor 555 (1:1000, Abcam Cat# ab150130; RRID:AB_2927775) and anti-rabbit Alexa Fluor 647 (1:500, Thermo Fisher Scientific Cat# A-31573; RRID:AB_2536183) for 1 hour at room temperature and DAPI (1:2000, Cell Signaling Cat# 4083). Finally, glomeruli were plated in 12% gelatin (Sigma-Aldrich, Cat# G7041) into glass bottom 35mm dishes (Greiner, Cat# 627860). Images were collected on a Leica TCS SP8 AOBS inverted confocal. When acquiring 3D optical stacks, confocal software was used to determine the optimal number of Z sections. Only the average intensity projections of these 3D stacks are shown in the results. Images were quantified using ImageJ.

#### In vivo – wild type and Alport mice

For *in vivo* experiments, AAV9-GFP was diluted in sterile buffer (6mM Tris hydroxymethyl aminomethane, 14mM Tris-HCl, 1mM MgCl^2^.6H^2^O, 0.2M NaCl, 0.005% P188, pH 8.0, Viralgen, Spain, Lot: 2023-10-10.01) on the day of injection to either low (1.32×10^12^ vg/mL), medium (6.6×10^12^ vg/mL), or high (3.3×10^13^ vg/mL) concentration and stored on ice until loaded into the syringe. AAV9-GFP was injected via the tail vein of the mice at 5mL/kg for low (6.60×10^12^ vg/kg), medium (3.30×10^13^ vg/kg), or high (1.65×10^14^ vg/kg) doses. Saline (0.9%) was injected via the tail vein for control groups. Administration of the AAV9-GFP was performed on 8-, 10-or 12-week-old WT and Col4a5 KO Alport mice (n=3-4 per group).

### Tissue collection, immunofluorescence and image quantification

At the defined study endpoint, four weeks post-injection, mice were humanely culled, and tissues (kidneys and liver) were collected and fixed in 4% paraformaldehyde (PFA) at 4°C for 48 hours. Fixed tissues were dehydrated, cleared, and infiltrated with paraffin wax before embedding into paraffin blocks. 5 µm sections were cut on a Leica RM2255 microtome and slides were left to dry overnight at 37°C. Antigen retrieval was performed by boiling tissue sections in citrate buffer (pH 6.0) for 20 min, followed by incubation at room temperature for 30 min. Samples were washed in PBS then blocked in 1% donkey serum, 2% BSA, and 0.1% Triton X-100 in PBS for 45 min at room temperature. Primary antibodies (rabbit anti-Wilms tumor protein WT1, 1:300, Abcam Cat# ab89901, RRID:AB_2043201; goat anti-GFP, 1:500, Cat# ab6673, RRID:AB_305643; rabbit anti-PDGFR beta, 1:100, Thermo Fisher Scientific Cat# MA5-15143, RRID:AB_10985851 and rat anti-CD31, 1:50, Dianova Cat# DIA-310-BA-2, RRID:AB_2916051) were diluted in blocking buffer and incubated overnight at 4°C in a humidified chamber. Secondary antibodies (Alexa Fluor 647 anti-rabbit; 1:400; Thermo Fisher Scientific Cat# A-31573; RRID:AB_2536183; Alexa Fluor 488 anti-rabbit; 1:400; Thermo Fisher Scientific Cat# A-21206; Alexa Fluor 647 anti-rat; 1:400; Thermo Fisher Scientific Cat# A-78947; Alexa Fluor 555 anti-goat; 1:400; Abcam Cat# ab150130; RRID:AB_2927775) were diluted in blocking buffer and incubated on slides for 45 min at room temperature in a humidified chamber. After washing, the slides were dried overnight and mounted with ProLongTM Gold Antifade mountant (ThermoFisher; P36930).

Whole-kidney and liver sections were imaged and stitched using the Olympus VS200 Slide-scanner (Olympus, Japan; RRID_SCR_024783) with a 20× objective in the DAPI, 555 and 647 channels. Images generated from slide scanning were viewed and fluorescence was quantified using QuPath (v.0.5.1; RRID:SCR_018257). 32 glomeruli per mouse were selected using the polygon tool and total cell number was determined using the cell detection feature in the DAPI channel. Single measurement classifiers were used to create thresholds for GFP and WT1 detection in the Cy3 and Cy5 channels, respectively. A composite classifier was created using the individual GFP and WT1 classifiers to quantify the total number of cells positive for GFP, WT1, or both markers. The object classifier was loaded onto all subsequent images. For higher resolution images Zeiss LSM 880 inverted AiryScan confocal was used with a Plan-Apochromat 40x/1.3 objective.

### Albumin-to-creatinine ratio

The albumin-to-creatinine ratio (ACR) test was performed as per the manufacturer instructions using the mouse albumin ELISA kit (Fortis Life science, E99-134) and creatinine kit (Bio-techne, KGE005).

### Flow cytometry

Glomeruli were isolated 4 weeks after AAV9-GFP administration to 10-week-old WT and Alport mice, and podocytes were labeled as previously described with slight modifications (20). Briefly, mice were perfused with 38 mL Hank’s balanced salt solution (HBSS) and 2 mL HBSS with enzymatic digestion solution (300U/mL Collagenase type II (Sigma), 50 U/mL DNase I (Invitrogen)) with 8×10^7^ Dynabeads™ M-450 Tosylactivated (Invitrogen). Kidneys were removed, minced into 1 mm^3^ pieces, and digested in 8mL enzymatic digestion buffer at 37°C for 20 min on a rotator. The digested kidneys were passed through a 100µm cell strainer and centrifuged at 200×g for 10 minutes at 4°C, and glomeruli were washed three times and collected using magnetic rack (Invitrogen). The purified glomeruli were digested by incubation in RPMI1640 (Sigma) containing 1 mg/ml Liberase TL (Roche), 100 U/ml DNase I, 10% FCS, 1% insulin/transferrin/selenium (Gibco), 1% penicillin/streptomycin, and 25 mM HEPES (Fisher Scientific) at 37°C for 2 hours with shaking at 1,400 rpm, and repeatedly mechanically sheared with a 27-gauge needle. After removing Dynabeads, cell suspensions were pelleted at 400×g at 4°C for 5 minutes. Subsequently, cells were blocked with FACS buffer (PBS with 0.5% BSA and 2 mM EDTA) containing normal syrian hamster serum and filtered 40µm strainer and then labelled with APC (allophycocyanin) conjugated anti-mouse Podoplanin Antibody (BioLegend, Cat# 127410; RRID:AB_10613649) at 4°C for 30 minutes in complete darkness. After washing twice with FACS buffer, cells were resuspended in FACS buffer containing DAPI solution (Thermo), analyzed by BD FACSymphony™ A5 and data processed using FlowJo v10.10 (RRID:SCR_008520).

### Statistical analysis

Statistical analysis was performed using GraphPad Prism (v9.3.1; RRID:SCR_002798). t-test or two-way ANOVA followed by Tukey’s multiple-comparison test were applied. Data are shown as mean ± SEM, with *P* values ≤0.05 (for t-test) and ≤ 0.01 (for two-way ANOVA) considered significant.

## Results

### Quantifying AAV9 transduction in podocytes in culture

We first investigated AAV9-GFP transduction in immortalized human podocytes (PODO/TERT) and isolated primary podocytes from wild-type and Alport (Col4a5 KO) mice. To evaluate the efficiency of transduction across these differentiated cell types, we performed a multiplicity of infection (MOI) assay, varying the viral genomes per cell from 5×10^4^ to 3.125×10^7^. We analyzed transduction of the GFP vector by immunofluorescence, quantifying the percentage of GFP-positive podocytes. At 24 hours post-transduction, we achieved 50% transduction in human PODO/TERT cells at a viral dose of 1.25×10^6^ vg/cell and 100% at 3.125×10^6^ vg/cell. We observed an equal transduction of both wild type and Alport primary mouse podocytes with 100% of GFP-positive cells at the MOI of 3.125×10^7^ vg/cell (**Figure 1A-B**). These results indicate that AAV9 effectively transduces podocytes in culture, with no difference observed between healthy and Alport podocytes.

**Figure 1:**
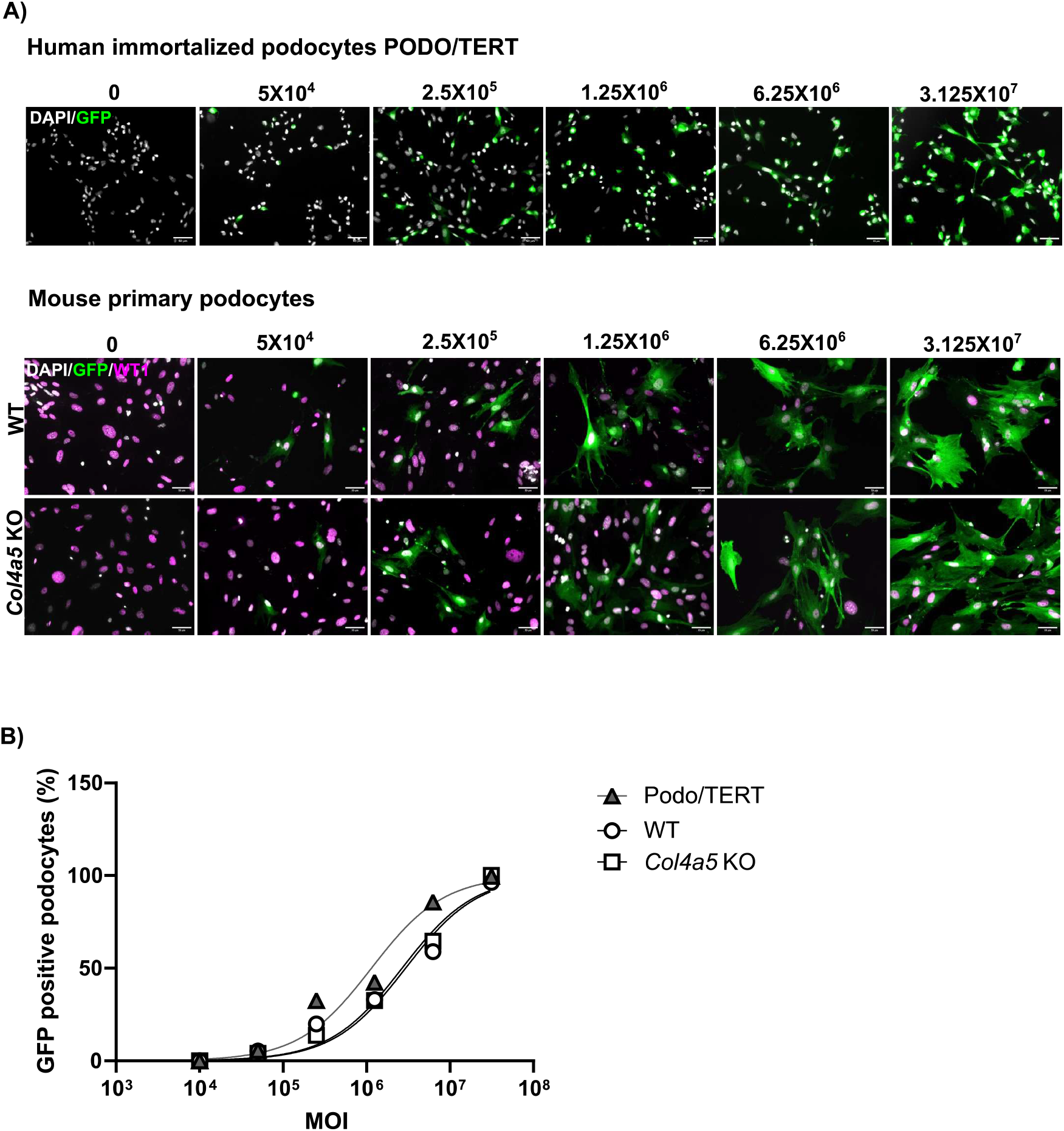
AAV9-GFP transduction of human immortalized and mouse primary podocytes. **A**) Human immortalized podocytes, PODO/TERT and mouse primary podocytes (wild type (WT) and Alport model Col4a5 KO) were transduced with AAV9-GFP at a multiplicity of infection (MOI) of 5×10^4^ to 3.125×10^7^ viral genomes (vg)/cell. The cells were analyzed by immunofluorescence at 24 hours post-transduction for the percentage of GFP positive cells. Scale bar = 50 µm. **B**) Estimation of percentage of cells infected based on MOI.

### Evaluating AAV9 transduction in ex-vivo glomeruli

Next, we established an *ex vivo* assay to evaluate whether AAV9-GFP could transduce isolated glomeruli from wild type and Alport (Col4a5 KO) mice. Isolated glomeruli from 8-week-old mice were incubated with 8.8×10^11^ viral genomes/mL of AAV9-GFP for 48 hours. We then evaluated glomerular GFP expression by immunofluorescence in transduced glomeruli from both wild-type and Alport mice and in untreated wild type glomeruli as control. The colocalization analysis between WT1, a podocyte marker, and GFP indicates that the GFP protein is expressed by podocytes in Alport *ex-vivo* glomeruli (**Figure 2A**). In addition, we observed a difference in the localization of GFP expression between the Alport and wild type glomeruli. The GFP signal was more peripheral in the wild type samples compared to Alport glomerulus in which the GFP viral particles appeared inside the glomeruli (**Figure 2A-B**). Of note, the Bowman’s capsule surrounding the glomeruli was kept intact in most of the wild-type samples during the isolation process but was absent in the Alport glomeruli possibly due to the loss of the collagen IV-alpha556 network (**S. Figure 2**). Quantification of the GFP signal revealed an increase by 2-fold of transduced cells inside the Alport glomeruli compared to wild type and a 4-fold increase of the peripheral signal in the wild type glomeruli compared to Alport (**Figure 2C**). Overall, the *ex-vivo* assay allows a comparison of AAV9-GFP transduction efficiency in glomeruli and demonstrates that an intact Bowman’s capsule limits the transduction of glomerular cells.

**Figure 2:**
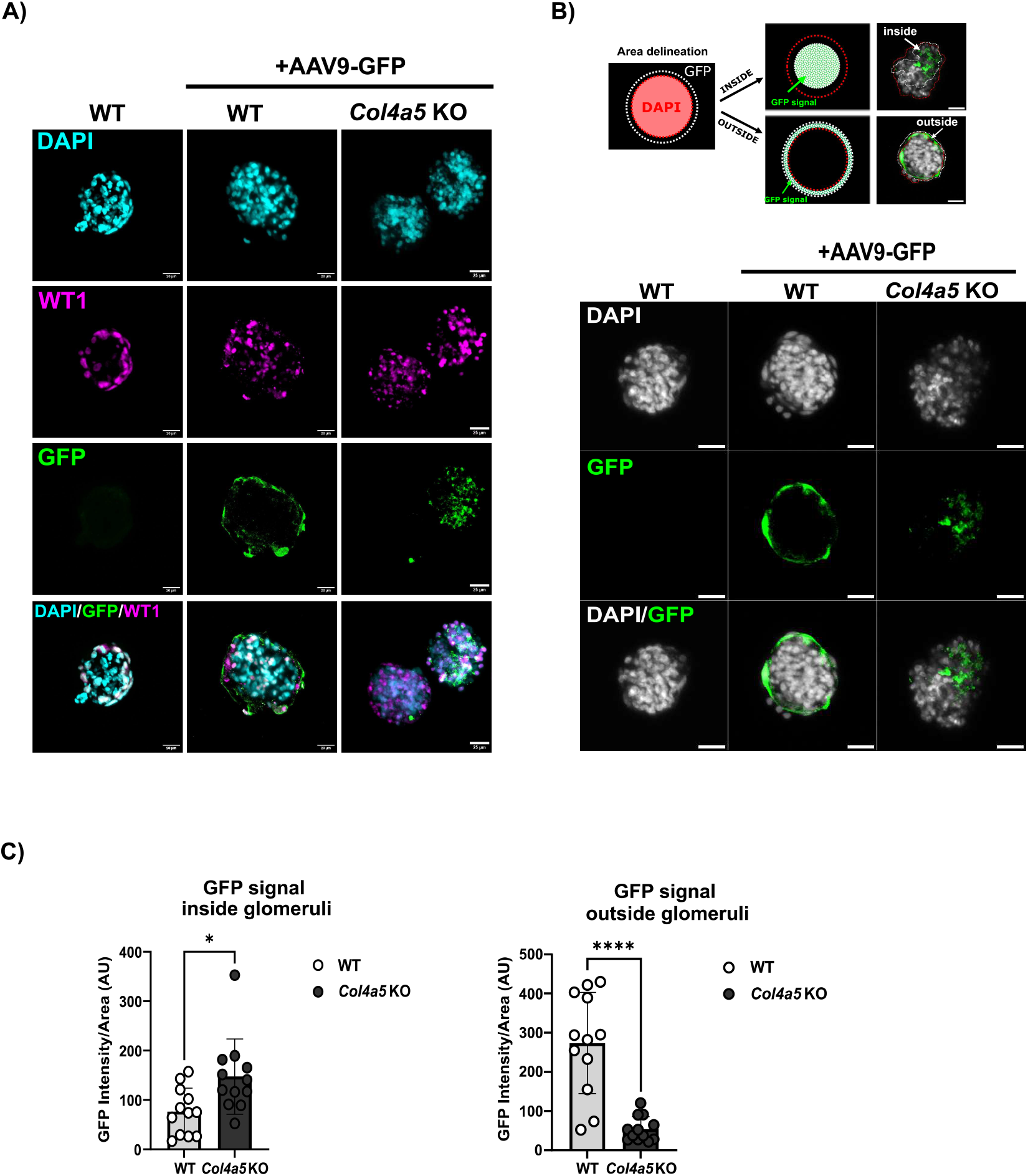
AAV9-GFP transduction of isolated mouse glomeruli. **A**) Isolated glomeruli (WT and Col4a5 KO) from 8-week-old mice were transduced with 8.8×1011 vg/ml of AAV9-GFP. GFP expression was visualized 48-hours post-transduction and co-stained with WT1, a podocyte marker. Scale bars = 25 µm. **B**) Schematic highlighting the area delineation used for measuring the GFP fluorescence intensity inside and outside the glomeruli. The red discontinued line represents the glomerulus, and the white line outlines the GFP signal. Representative images illustrating the localization of the GFP signal in WT and Alport ex-vivo glomeruli. Scale bars = 25 µm. **C**) Quantification of the GFP fluorescence intensity inside and outside of WT and Col4a5 KO Alport glomeruli (n=12 per genotype). *P* value <0.0001 (***), p value<0.01 (*).

### Increased AAV9 transduction in Alport podocytes

Finally, we examined the transduction efficiency of AAV9-GFP administered to mice via tail-vein injection. We tested low, medium, and high doses of AAV9-GFP (6.60×10^12^, 3.30×10^13^, 1.65×10^14^ vg/kg respectively) in 8-week-old WT and Alport mice and harvested tissues four weeks post-injection (**Figure 3A**). Two cross-sections per whole kidney were probed for the GFP transgene and the podocyte cell marker WT1 (**Figure 3B**). We then quantified GFP-positive cells in the kidney and in the liver. We found 10-20% GFP-positive cells in the liver of WT and Alport mice across all doses, demonstrating successful delivery of the AAV9-GFP. Liver sections were further assessed by picrosirius red staining as an indicator of fibrosis, but no differences were detected between control and AAV-injected mice (**S. Figure 3**). In the kidney, we found no significant difference in AAV9-GFP transduction efficiency between healthy and Alport mice at low and medium doses (**Figure 3C-D**). In the mice injected with high dose, we detected GFP protein in 10% of Alport podocytes compared to 3% in the wild type (**Figure 3E**). Notably, we detected variability in the uptake of AAV9-GFP in glomeruli within individual animals, with a proportion of glomeruli negative for the GFP vector. High-resolution imaging showed that at the medium dose, podocytes start to express GFP in Alport mice and at the high dose, a greater number of podocytes were transduced in Alport compared to WT mice (**Figure 3D**, white arrow).

**Figure 3:**
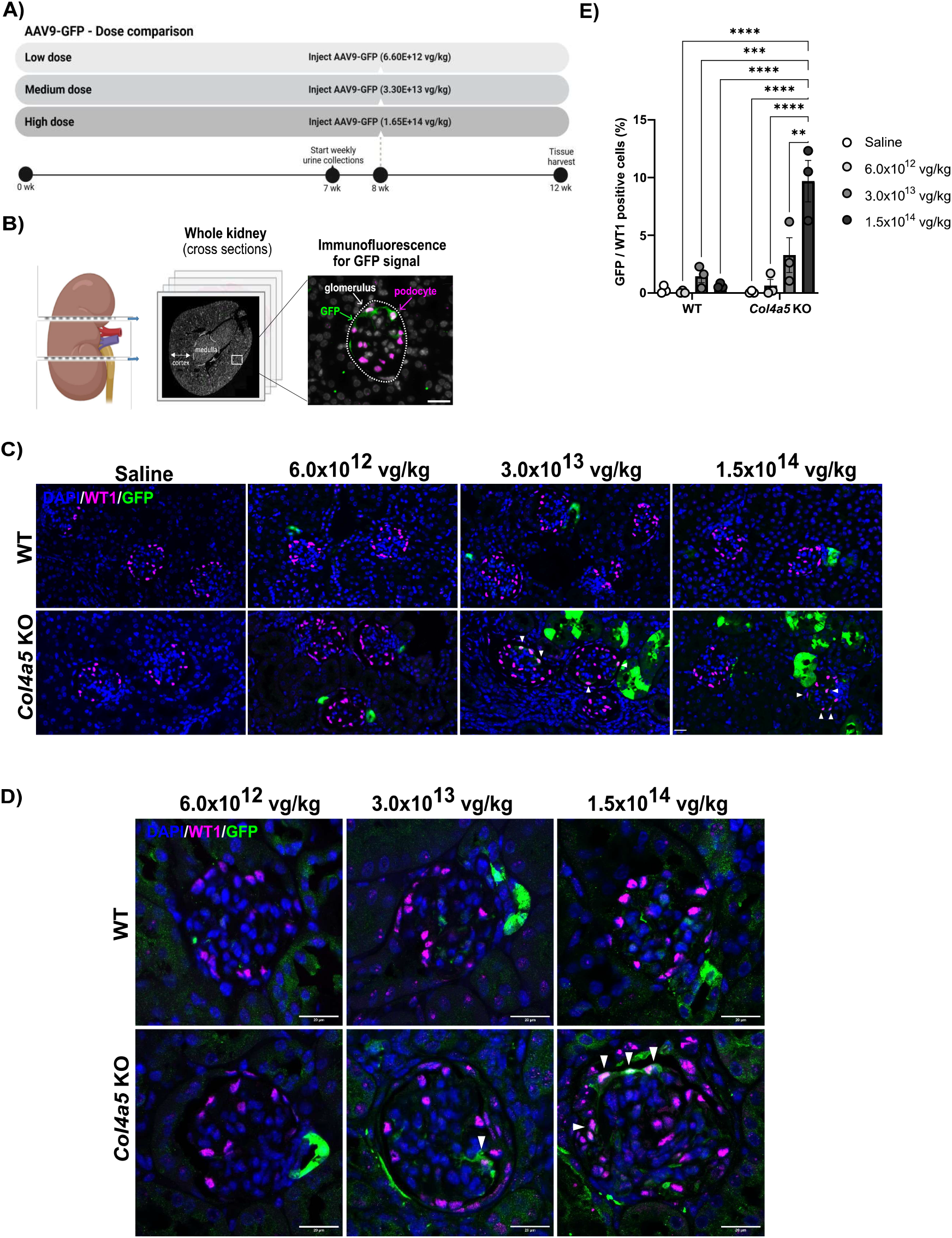
AAV9-GFP transduction in kidneys of wild type and Alport mice injected at low (6×10^12^ vg/kg), medium (3.0×10^13^ vg/kg) and high (1.5×10^14^ vg/kg) dose. In total, 6 (3 WT and 3 Col4a5 KO) 8-week-old mice per group were injected with either AAV9-GFP or saline. **A**) Study design for the AAV9-GFP dose comparison. **B**) Schematic demonstrating the workflow for image analysis. 2 kidney cross-sections were used for quantification where all glomeruli were identified by WT1 immunofluorescence (podocytes, magenta) and GFP-positive podocytes were counted (green). **C**) Representative immunofluorescence images of glomerular sections where DAPI is in grey, WT1 (podocyte marker) in magenta and GFP in green. Scale bars = 20 µm. **D**) High-resolution images showing the GFP localization in WT and Alport glomeruli from mice injected with the three AAV9-GFP doses. White arrows highlight colocalization of the GFP vector with the podocyte marker. **E**) Colocalization for WT1 and GFP signals was quantified using QuPath and plotted as the percentage of GFP-positive podocytes. The entirety of glomeruli from 2 whole kidney cross-sections (per genotype/condition) were quantified. Data are presented as mean ± SEM of the values from 3 mice per group and per genotype. P value<0.05 (*), <0.005 (**), <0.0005 (***).

Next, we assessed AAV9-GFP transduction during kidney disease progression, by injecting 8-, 10-, and 12-week-old WT and Alport mice via tail vein with a high dose of the virus (**Figure 4A**). To monitor disease progression and a possible effect of viral transduction we measured albumin:creatinine ratios (ACR). As expected, we observed an increase in ACR with age in Alport mice but importantly, there were no difference between saline and AAV9-GFP treated mice (**Figure 4B**). Immunofluorescence revealed that the number of GFP-positive podocytes increased by 4-fold in the Alport mice injected at 12-week-old compared to 8-week-old, reaching around 20% of the cells (**Figure 4C-E**), indicating that Alport podocytes are more readily transduced at the later stage of the disease. We also observed an increased GFP signal in the tubules as the disease progressed (**Figure 4C-D**).

**Figure 4:**
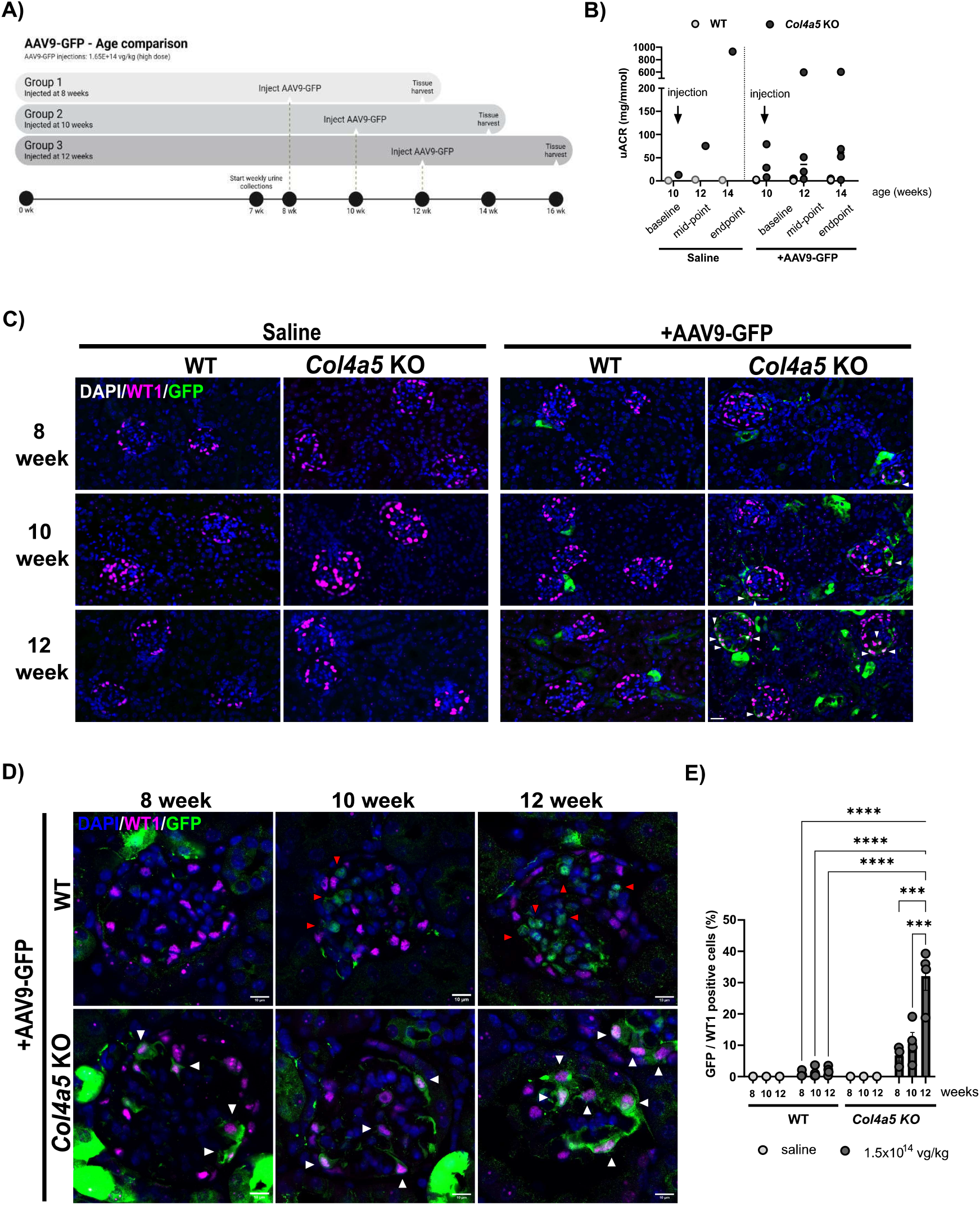
AAV9-GFP transduction in kidneys of 8, 10, 12-week-old wild type and Alport mice injected with saline or the high dose (1.5 × 10^14^ vg/kg). **A**) Study design for the AAV9-GFP age comparison. **B**) Urinary ACR performed on 10-week-old mice (WT and Alport) after injection of either saline or high-dose of AAV9-GFP. Urine was collected for baseline read-out at 10 weeks (before injection), mid-point at 12 weeks (2 weeks post-injection) and endpoint at 14 weeks (4 weeks post-injection). **C**) Representative immunofluorescence images of glomerular sections from tissue collected 4 weeks post-injection and stained with DAPI, WT1 and GFP. Scale bars = 20 µm. **D**) High-resolution images showing the GFP localization in WT and Alport glomeruli from mice injected with the high dose of AAV9-GFP. White arrows highlight colocalization of the GFP vector within the podocytes and red arrows, the localization of GFP with other glomerular cells. **E**) The quantification of GFP-positive podocytes (%) is displayed for the two phenotypes WT (left) and Alport mice (right), injected with either saline (light grey bars) or high dose of AVV9-GFP (dark grey bars) (n=2-4 mice per group and per condition). Data are presented as mean ± SEM. *P* value <0.005 (**).

These findings were supported by flow cytometry analysis. Isolated primary glomerular cells from 10-week-old WT and Alport mice were immunolabelled to detect podoplanin, a podocyte marker and the number of podoplanin-GFP double positive cells were quantified. Between 8-17% of Alport podocytes were GFP positive, compared to 4-6% of WT cells **(Figure 5A-B)**. Furthermore, the Alport mice displayed a lower number of podocytes compared to wild type, but a higher percentage of GFP positive podocytes indicating that AAV9-GFP is more likely to reach podocytes in injured glomeruli (**Figure 5C**).

**Figure 5:**
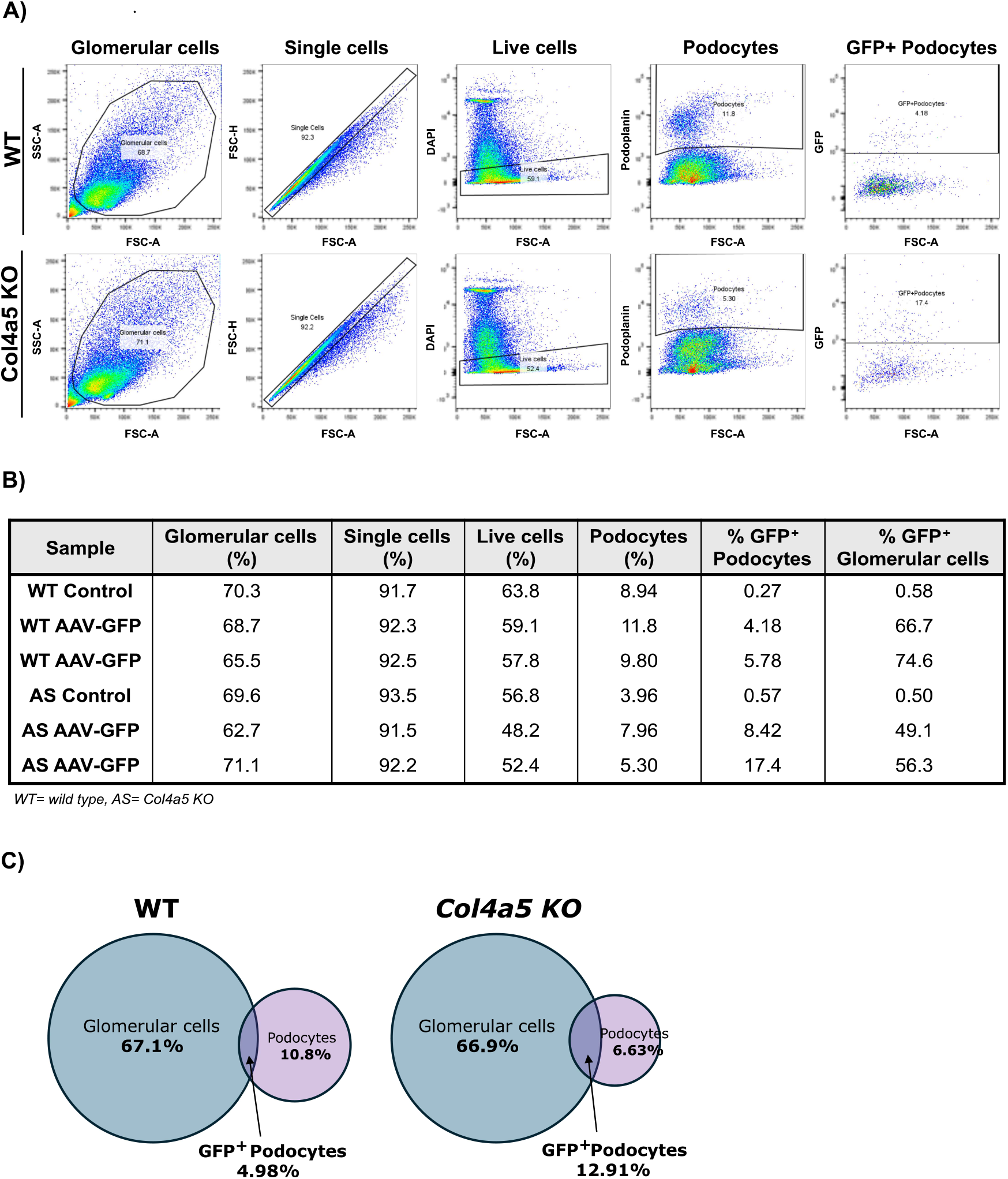
Flow cytometry analysis of single cells isolated from glomeruli, 4 weeks after AAV-GFP administration to 10-week-old WT and Alport mice. **A**) Flow cytometry gating strategy and representative scatterplots. After gating glomerular single cells, DAPI negative cells were selected (viable cells) and, within those, podoplanin positive cells (podocytes). Analysis of GFP gate represents the transduced podocytes. **B**) Data showing the percentage of glomerular cells, single cells, live cells, podocytes and GFP-positive podocytes in WT and Alport (AS) (n=2 WT+AAV-GFP, n=1 WT control saline; n=2 Col4a5 KO (AS)+AAV-GFP, n=1 Col4a5 KO control saline). **C**) Venn diagrams representing the proportion of isolated glomerular cells, podocytes and GFP-positive podocytes in WT and Alport mice.

Interestingly, we observed that in Alport mice, the GFP transgene was predominantly colocalized in podocytes, particularly as the disease progresses, however in WT mice, the GFP is expressed in other glomerular cell types (**Figures 4D**, red arrows). To determine which cells were transduced by AAV9-GFP, kidney sections were immunolabelled with mesangial and endothelial markers (PDGFR beta and CD31 respectively). In WT mice, the GFP vector mainly colocalized with PDGFR-positive mesangial cells and not with endothelial cells (**Figure 6**).

**Figure 6:**
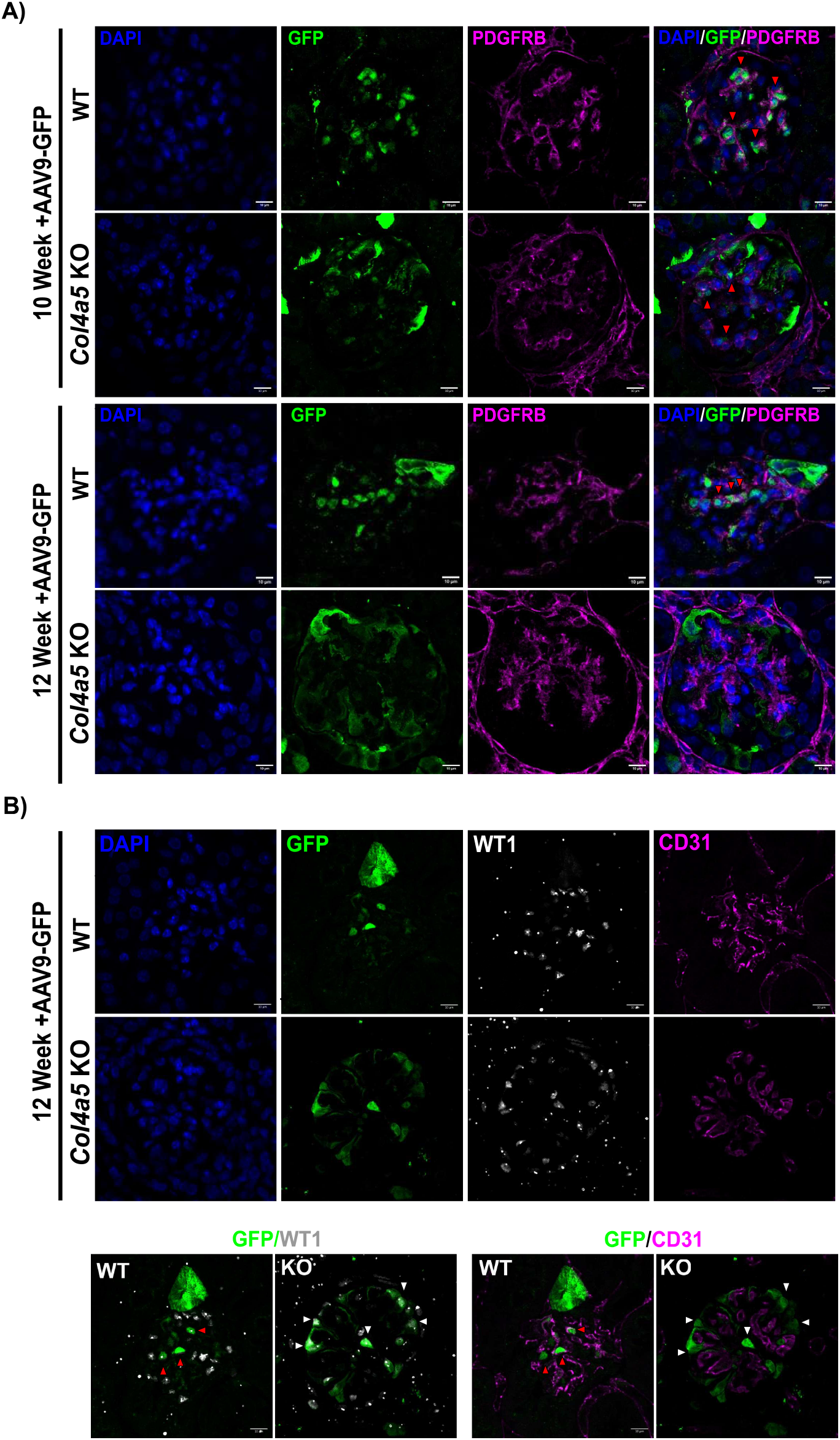
Immunofluorescence detection of AA9-GFP in mesangial and endothelial cells. **A**) High-resolution images showing the GFP localization in WT and Alport glomeruli from mice injected with high dose of AAV9-GFP at 10 and 12 weeks. Glomerular sections stained with a mesangial marker, PDGFRB. Red arrows highlight the expression of the GFP in mesangial cells. **B**) Representative images of glomeruli from WT and Alport mice injected with high dose of AAV9-GFP at 12 weeks. Kidney sections stained with WT1 and CD31, an endothelial marker. Scale bars = 20 µm.

Overall, these data demonstrate that AAV9-GFP effectively transduces podocytes, with enhanced delivery in a dose-dependent manner and increased efficiency in mice with more compromised glomerular basement membranes at a later stage of kidney disease progression.

## Discussion

This AAV9-GFP proof-of-concept study highlights gene therapy as a promising therapeutic strategy for monogenic kidney diseases such as Alport syndrome. We found that AAV9-GFP can reach the glomerular filtration barrier in the kidneys of injected mice and efficiently transduce podocytes. Notably, we observed that there was enhanced delivery of the transgene in podocytes in an Alport model compared to healthy control. Additionally, the transduction efficiency increased in a dose-dependent manner and with disease progression.

The increased permeability of the glomerular filtration barrier in Alport syndrome is hypothesized to contribute to the enhanced AAV9 transduction. This concept of the altered structure of the glomerular basement membrane facilitating the penetration of AAV9 was previously reported in a CKD and Alport model (16, 21). In these models, both podocytes and tubular cells were transduced, a finding which we also observed. In our Col4a5 KO Alport model, the podocyte uptake of AAV9 was significantly reduced in mice injected at earlier time points (8 and 10-week-old) when the disease was less established, while the highest GFP transgene expression was seen at 12 weeks. Transduction of mesangial cells by AAV was also previously reported (14). In our study, the WT mice injected at 10 and 12 weeks showed that AAV9-GFP localized primarily to mesangial cells, and this was also observed to a lesser extent in the Alport mice. This suggests that the stage of disease at the time of injection plays a critical role in determining the efficiency of AAV transduction.

Despite these promising findings, challenges remain, particularly regarding the effective delivery of AAV9 to target tissues. The choice of administration route can significantly impact therapeutic outcomes. One possible improvement to reduce side effects on other organs could be using different delivery routes, such as direct injection into the renal artery instead of a peripheral intravenous injection (22, 23). Renal vein injections have also been found to be effective for kidney-targeted gene delivery in healthy mice, achieving high transduction efficiency in the cortex and medulla, targeting both glomeruli and tubules (23). However, we have shown that in an Alport model, AAV9 injected via a peripheral intravenous route was preferentially distributed to podocytes, indicating that this route of administration may be sufficient with certain forms of genetic kidney disease.

To further enhance transduction efficiency, and cell specificity additional strategies could be employed, including tailoring the AAV by customizing cell-specific promoters, optimizing transgene designs, and engineering capsid variants to achieve specific tissue tropism (24).

Whilst the findings of this and other studies is encouraging, there are additional hurdles to overcome for gene therapy in Alport syndrome. These include the limited cargo capacity of AAV, the COL4 genes are too large to fit within the AAV vector, as they are approximately 5.05 kilobases (kb) in size and AAV has limited packaging capacity, considered to be around 4.7-5 kb (25). Podocyte secretion of transduced collagen IV, and subsequent assembly into the basement membrane also need to be overcome. Finally, we do not yet know the level of transduction efficiency required for meaningful rescue of Alport type IV collagen deficiency and if this attainable (10). Nevertheless, our findings of increased transduction and expression of AAV9-GFP in Alport podocytes demonstrate that effective therapies may be possible using AAV vectors.

## Data availability

### Extended data

This project contains the following underlying data:

-Extended data figures.pdf:

S. Figure 1-AAV9-GFP-WPRE reporter plasmid

S. Figure 2 - Images panel of isolated mouse glomeruli transduced with AAV9-GFP

S. Figure 3 - Representative images of liver cross-sections from 12-week-old mice injected with AAV9-GFP

## Author Contributions

R.L., D.R.S, A.S and K.G, conceptualization; E.W, G.B, M.F., S.K., investigation; R.L., M.F, A.A., G.B. and E.W. writing original draft; R.L., D.R.S, A.S, K.G., E.W., G.B., A.A, S.K., M.F., reviewing and editing manuscript.

## Competing interests

The authors declare the following conflicts of interest:

R.L., D.R.S, E.W., M.F., G.B., A.A., A.S. and K.G. work on a collaborative research project funded by FourPoints Innovation, a portfolio company of certain funds managed by Deerfield Management Company, L.P.

R.L. has the following additional disclosures:

Clinical trials PI: Bayer, Travere, River3Renal, Calliditas.

Collaborative research agreements: Calliditas, Novo Nordisk.

## Grant information

R.L. is funded by the Wellcome Trust (Ref: 226804/Z/22/Z, Ref: 227417/Z/23/Z, Ref: 301803/Z/23/Z), Kidney Research UK and The Stoneygate Trust (Alport Research Hub) and the National Institute for Health Research (NIHR-Manchester BRC). S.K is funded through a Japan Agency for Medical Research and Development (AMED) fellowship.

## Acknowledgements

We thank the University of Manchester Biological Support Facility for assistance in animal welfare, the Bioimaging Core Facility and the Flow Cytometry Core Facility. We thank Conor Sugden for advice with image analysis.

