## Supplementary materials for "Increased *in vivo* transduction of AAV-9 cargo in Alport podocytes"

### Supplementary Figure 1

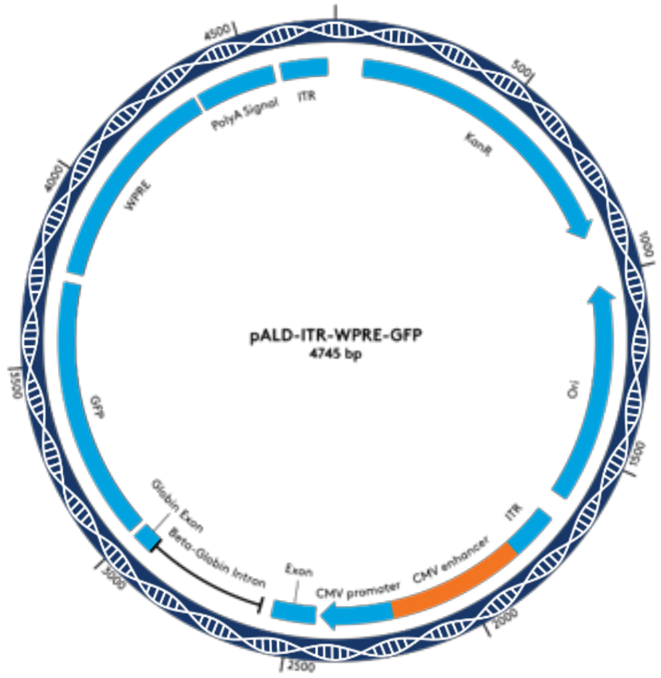


S. Figure 1: rAAV9-GFP-WPRE reporter plasmid. Circular map of the AAV9-GFP reporter construct with the regulatory elements.

#### Supplementary Figure 2

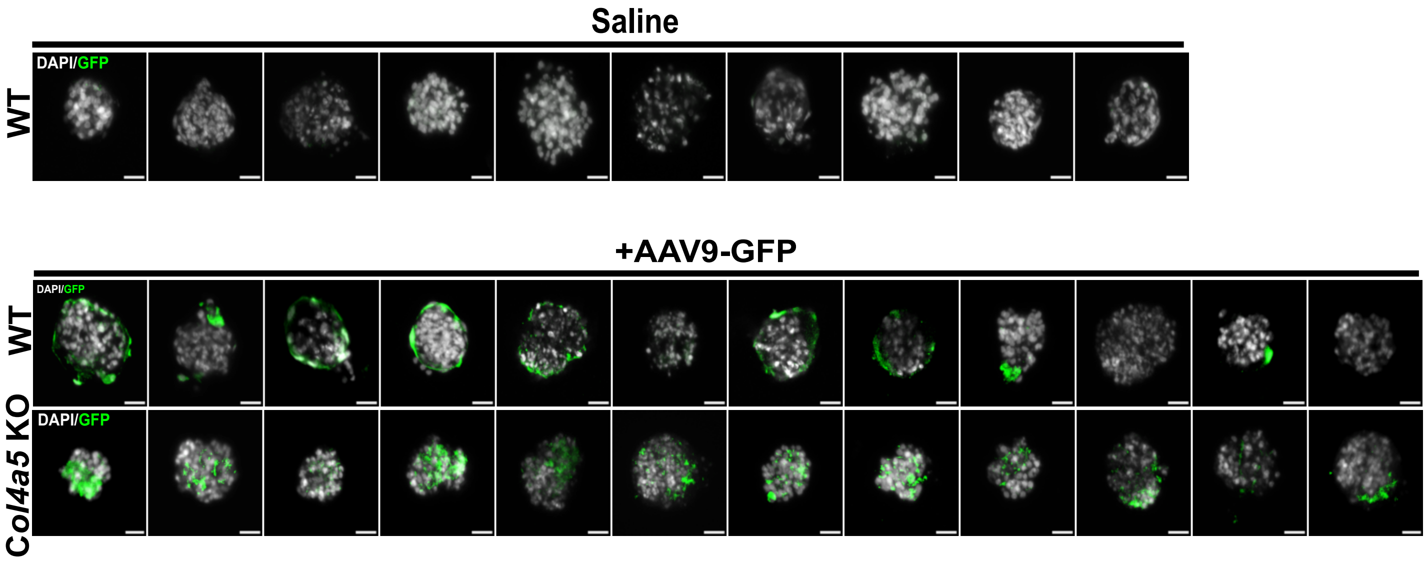


S. Figure 2: Panel images of isolated mouse glomeruli transduced with rAAV9-GFP. Images of 10 wild type glomeruli isolated from 8-week-old mice (control, no rAAV9-GFP); 12 wild type and Col4a5 KO glomeruli transduced with rAAV9-GFP. Scale bar= 25 µm.

### Supplementary Figure 3

A)

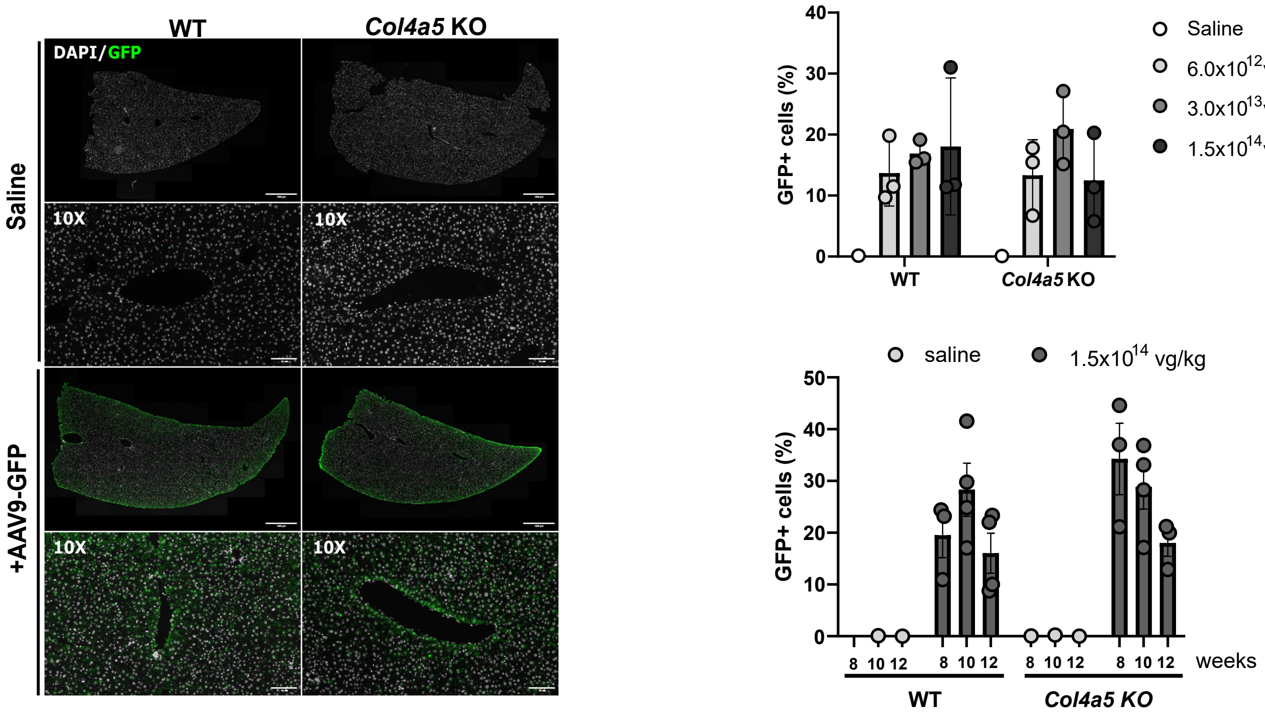


B)

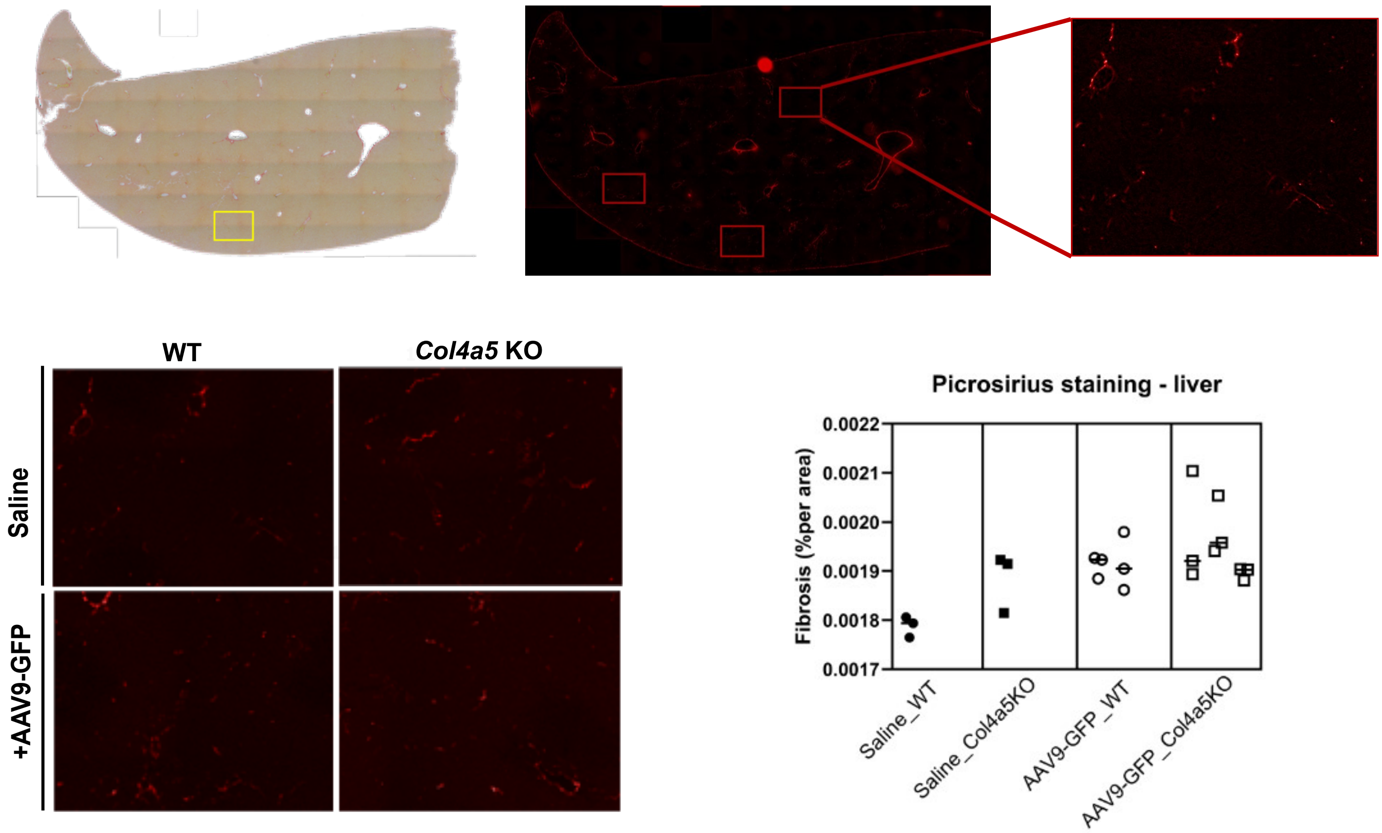


S.Figure 3: Images of whole liver of wild type and Alport mouse injected with rAAV9-GFP. A) Representative images of liver cross-sections from 12-week-old mice injected with 1.5 x 1014 vg/kg (high dose) of rAAV9-GFP. Quantification of GFP intensity in the liver. Scale bar= 1000 µm. Zoomed in images, scale bar= 100 µm. B) Schematic representation and quantification of liver stained with picrosirius red.

**Method:**

*Picrosirius staining and quantification pipeline*

Slides were dewaxed using a Leica Autostainer XL CV5030 then stained using Picrosirius red solution for 1 hour and washed with 1% acetic acid. Slides were dehydrated, cleared and mounted using the Leica Autostainer XL CV5030.

Liver sections were imaged by an Olympus BX63 upright microscope using a 20x objective using colour mode in CellSens Dimension v1.16. Images were stitched together to create whole section images and then processed in ImageJ. To quantify collagen and fibrosis, the rectangle tool was used to select 3 uniform areas per liver section, excluding regions containing vessels or ducts. Images were split into colour channels and the red channel was used to measure mean grey intensity. Collagen density was measured by dividing mean grey intensity by surface area.
